# Influence of mutagenic versus non-mutagenic pre-operative chemotherapy on the immune infiltration of breast cancer

**DOI:** 10.1101/455055

**Authors:** Anna-Mária Tőkés, Orsolya Rusz, Gábor Cserni, Erika Tóth, Gábor Rubovszky, Tímea Tőkés, Laura Vízkeleti, Lilla Reiniger, Renáta Kószó, Zsuzsanna Kahán, Janina Kulka, Marco Donia, András Vörös, Zoltan Szallasi

**Affiliations:** 2^nd^ Department of Pathology, Semmelweis University, Budapest, Hungary; Department of Oncotherapy, University of Szeged, Szeged, Hungary; Department of Pathology, University of Szeged, Szeged, Hungary; Department of Pathology, Bács-Kiskun County Teaching Hospital, Kecskemét, Hungary; National Institute of Oncology, Budapest, Hungary; Oncology Center, Semmelweis University, Budapest, Hungary; MTA-SE-NAP B Brain Metastasis Research Group, 2^nd^ Department of Pathology, Semmelweis University, Budapest, Hungary.; 1^st^ Department of Pathology and Experimental Cancer Research, Semmelweis University, Budapest, Hungary; Center for Cancer Immune Therapy, Department of Hematology, Herlev Hospital, University of Copenhagen, Copenhagen, Denmark; Department of Oncology, Herlev Hospital, University of Copenhagen, Copenhagen, Denmark; Department of Bio and Health Informatics, Technical University of Denmark, Lyngby, Denmark; Computational Health Informatics Program, Boston Children’s Hospital, Harvard Medical School, Boston, USA

**Keywords:** Tumor infiltrating lymphocytes, mutagenic capacity, neoadjuvant treatment, breast cancer

## Abstract

**Background:** Chemotherapeutic agents are often mutagenic. Induction of mutation associated neo-epitopes is one of the mechanisms by which chemotherapy is thought to increase the number of tumor-infiltrating lymphocytes, but the clinical relevance of this triggered immune response is not known.

We decided to investigate, whether treatment with various chemotherapeutic agents with significantly different mutagenic capacity induce a significantly different number of stromal tumor-infiltrating lymphocytes (StrTIL) in the clinical setting.

**Methods:** 112 breast carcinoma cases treated with pre-operative chemotherapy were selected for the study. According to chemotherapy regimen 28/112 patients received platinum-based, 42/112 cyclophosphamide-based and 42/112 anthracycline-based chemotherapy. The percentage of stromal tumor-infiltrating lymphocytes (StrTIL) was evaluated on hematoxylin-eosin stained slides of pre-treatment core biopsy (pre-StrTIL) and post-treatment surgical tumor samples (post-StrTIL), according to the most recent recommendation of International TILs Working Group. In survival analyses, TIL changes (ΔStrTIL) were calculated from the difference between post-StrTIL and pre-StrTIL.

**Results:** Of the 112 cases, 58.0% (n=65) were hormone receptor (HR) positive and 42.0% (n=47) were HR negative. There was a trend of higher post-StrTIL compared to pre-StrTIL (median 6.25% vs. 3.00%; p<0.001). When analyzing the pre-StrTIL and post-StrTIL among the three treatment groups, we experienced significant StrTIL increase independently of the treatment applied. Based on the results of survival analyses both post-StrTIL and ΔStrTIL was found to be independent prognostic factor in HR negative cases. Each 1% increase in post-StrTIL reduced the hazard of distant metastases development by 2.6% (hazard ratio: 0.974; CI: 0.948-1.000; p=0.05) and for each 1% ΔStrTIL increment, the risk of distant metastases was reduced by 4.3% (hazard ratio: 0.957; CI: 0.932-0.983; p=0.001). The prognostic role of StrTIL in HR positive cases could not be proven.

**Conclusions:** StrTIL expression might be stimulated by highly (platinum), moderately (cyclophosphamide) and marginally (taxane, anthracycline) mutagenic chemotherapeutic agents. Increase in StrTIL in residual cancer compared to pre-treatment tumor tissue is associated with improved distant metastasis-free survival in cases with HR negative breast carcinoma.

## Background

Appropriate stimulation of a patient’s immune response can result in durable control of widely metastatic solid tumors. However, such clear clinical benefit is observed only in a relatively small proportion of patients [1,2].

An increasing body of experimental evidence underlines the importance of somatic coding genetic alterations (i.e. mutations) for recognition of cancer by the human immune system. The various forms of coding mutations are often translated into altered proteins including novel peptide sequences, which can become “neo-epitopes” on the surface of tumor cells ready to be scanned by the patient’s immune repertoire of T cells. These neo-epitopes are believed to be particularly immunogenic because they are not encoded in the normal genome of the individual patient, thus reactive T cells are not subjected to central tolerance. Recognition of neo-epitopes by cytotoxic T cells can lead to immune-mediated tumor regression. However, given the highly variable and individual nature of somatic tumor mutations, specific interventions to target neo-epitopes are technically very challenging.

The burden of neo-epitopes (which is strongly correlated with the total mutational load) is a strong predictor of response to current cancer immunotherapies [3,4]. This association is thought to reflect the level of how “foreign” is a given tumor, which would then be associated with stronger immune responses.

We have previously experimentally classified current chemotherapy regimens as highly (cisplatin), moderately (cyclophosphamide) and marginally/non (paclitaxel, doxorubicin and gemcitabine) mutagenic. We found that platinum therapy induces 20-fold and cyclophosphamide therapy induces 5-fold more mutations and neo-epitopes than the number of neo-epitopes present in taxane or anthracycline treated tumors [5]. Since the average rate of mutations for breast cancer is about 1 mutation/Megabase, a 20-fold increase of mutations would shift the average rate of mutations to the level seen in melanoma [6]. Building on these data, we hypothesized that induction of a higher number of mutations and neo-epitopes with mutagenic chemotherapy might result in stronger immune reactions in the tumor microenvironment, and this could be reflected by a larger increase in lymphocytic infiltration. This prompted us to investigate whether highly mutagenic chemotherapy induces a larger increase in lymphocytic infiltration compared to low/non-mutagenic chemotherapy when administered as pre-operative treatment for breast cancer.

## Methods

### Patients

112 patients diagnosed with breast carcinoma and treated with pre-operative chemotherapy in four Hungarian institutions (National Institute of Oncology, Onco-Radiology Center of Bács-Kiskun County Teaching Hospital, Semmelweis University and University of Szeged) between 2005 and 2017, were selected and their samples studied retrospectively. The inclusion criteria were as follows: availability of both core biopsy and surgical tumor sample, known clinical and treatment data, at least 2 cycles of chemotherapy administered before surgery, residual tumor after pre-operative chemotherapy. All patients underwent breast surgery. Of the 112 cases, 103 received chemotherapy plus surgery with curative intent, while 9 cases had bone metastases at the beginning of pre-operative chemotherapy. The tumor histological type was defined according to the most recent World Health Organization’s classification [6].

Hormone receptor (HR) status was scored according to the respective current Hungarian Guidelines [7] and the American Society of Clinical Oncology/College of American Pathologists’ recommendations [8]. A case was considered HR negative if the expression of estrogen receptor and progesterone receptor was less than 1%. HER2 positivity was evaluated conforming to the United Kingdom recommendations [9]. Based on the HR and HER2 statuses, cases were grouped into four different subtypes (Table 1)

**Table 1.**
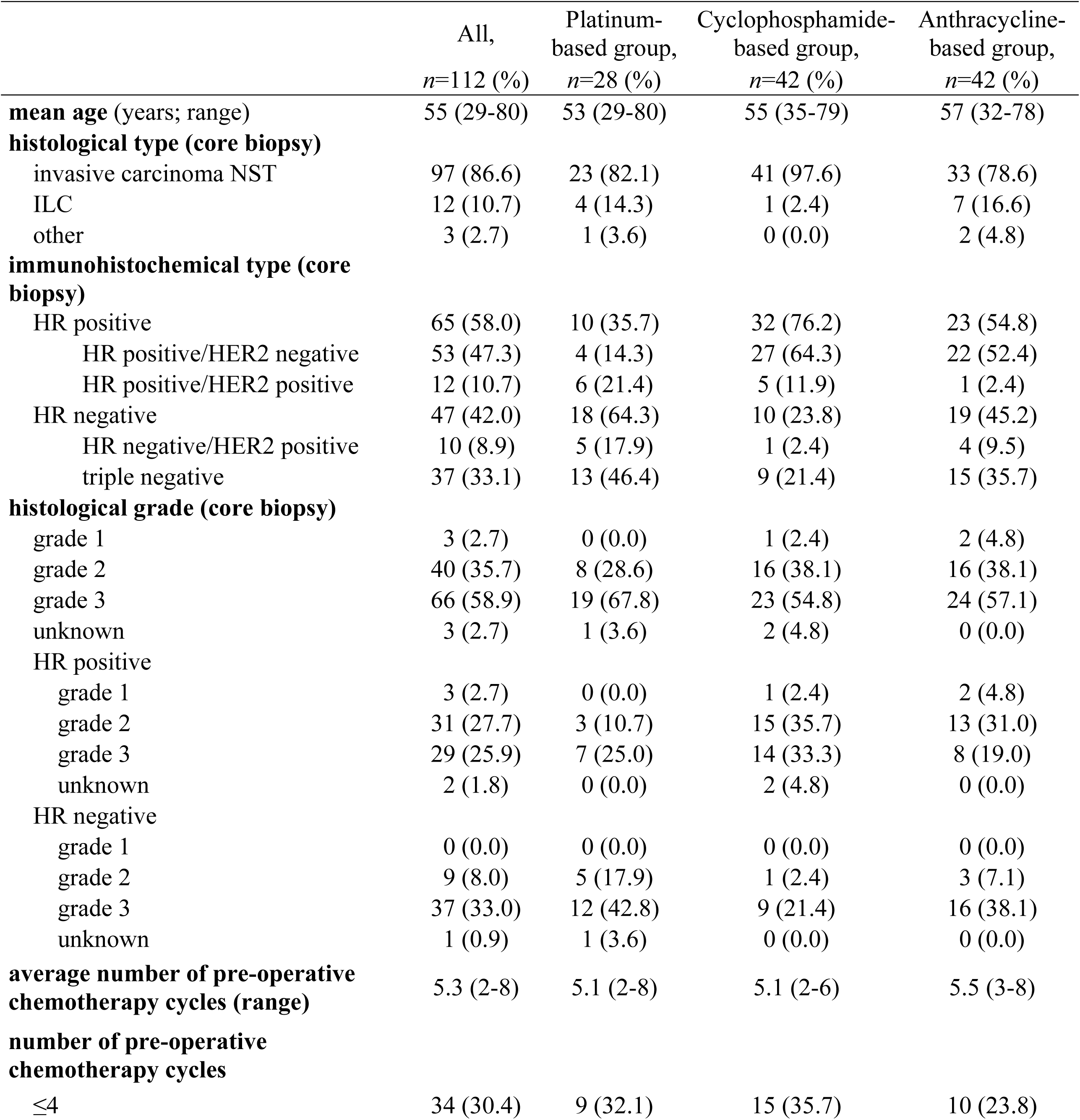
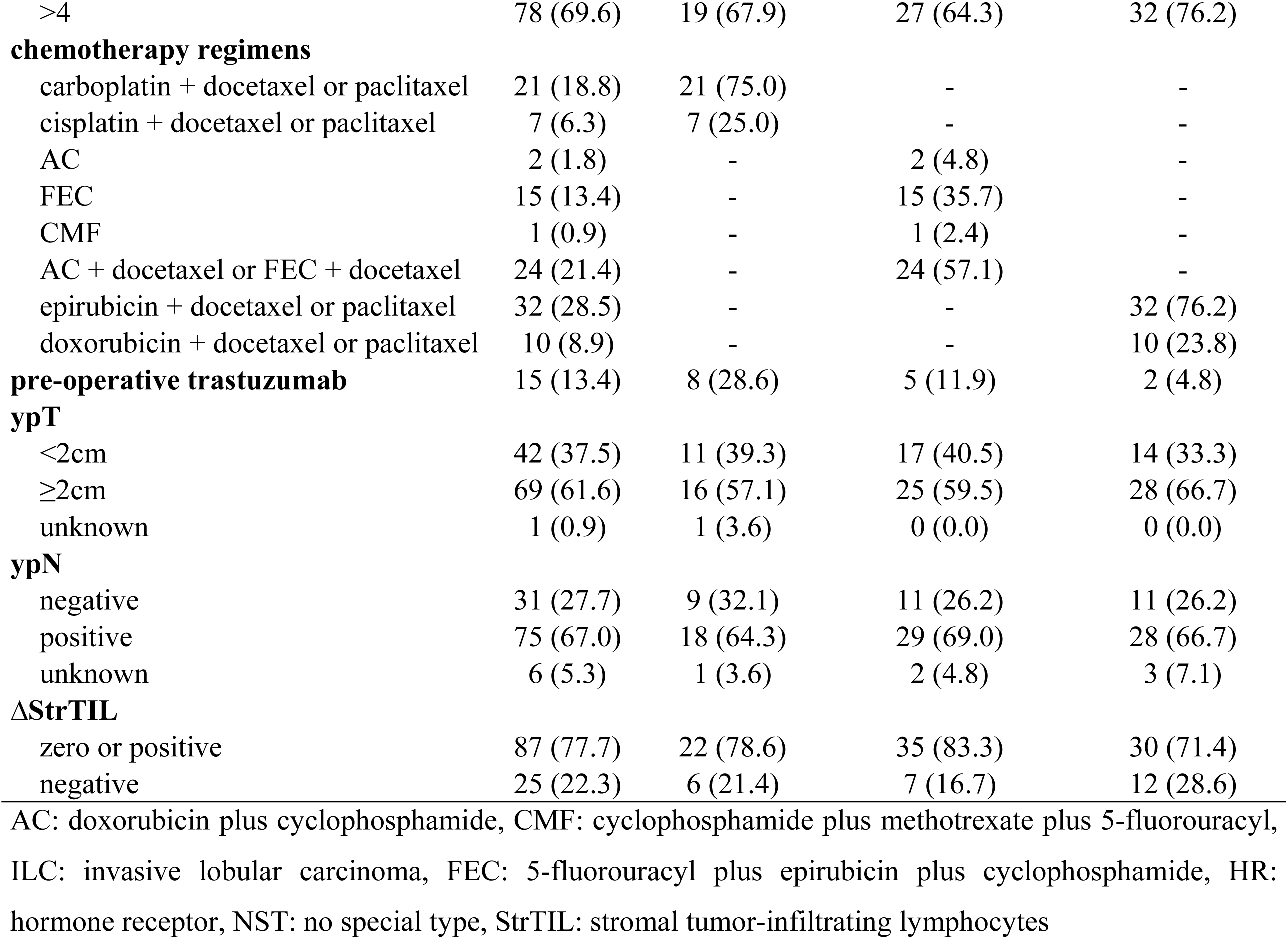
Clinico-pathological characteristics

|  | All,<br>*n*=112 (%) | Platinum-based group,<br>*n*=28 (%) | Cyclophosphamide-based group,<br>*n*=42 (%) | Anthracycline-based group,<br>*n*=42 (%) |
| --- | --- | --- | --- | --- |
| **mean age** (years; range) | 55 (29-80) | 53 (29-80) | 55 (35-79) | 57 (32-78) |
| **histological type (core biopsy)** |  |  |  |  |
| invasive carcinoma NST | 97 (86.6) | 23 (82.1) | 41 (97.6) | 33 (78.6) |
| ILC | 12 (10.7) | 4 (14.3) | 1 (2.4) | 7 (16.6) |
| other | 3 (2.7) | 1 (3.6) | 0 (0.0) | 2 (4.8) |
| **immunohistochemical type (core biopsy)** |  |  |  |  |
| HR positive | 65 (58.0) | 10 (35.7) | 32 (76.2) | 23 (54.8) |
| HR positive/HER2 negative | 53 (47.3) | 4 (14.3) | 27 (64.3) | 22 (52.4) |
| HR positive/HER2 positive | 12 (10.7) | 6 (21.4) | 5 (11.9) | 1 (2.4) |
| HR negative | 47 (42.0) | 18 (64.3) | 10 (23.8) | 19 (45.2) |
| HR negative/HER2 positive | 10 (8.9) | 5 (17.9) | 1 (2.4) | 4 (9.5) |
| triple negative | 37 (33.1) | 13 (46.4) | 9 (21.4) | 15 (35.7) |
| **histological grade (core biopsy)** |  |  |  |  |
| grade 1 | 3 (2.7) | 0 (0.0) | 1 (2.4) | 2 (4.8) |
| grade 2 | 40 (35.7) | 8 (28.6) | 16 (38.1) | 16 (38.1) |
| grade 3 | 66 (58.9) | 19 (67.8) | 23 (54.8) | 24 (57.1) |
| unknown | 3 (2.7) | 1 (3.6) | 2 (4.8) | 0 (0.0) |
| HR positive |  |  |  |  |
| grade 1 | 3 (2.7) | 0 (0.0) | 1 (2.4) | 2 (4.8) |
| grade 2 | 31 (27.7) | 3 (10.7) | 15 (35.7) | 13 (31.0) |
| grade 3 | 29 (25.9) | 7 (25.0) | 14 (33.3) | 8 (19.0) |
| unknown | 2 (1.8) | 0 (0.0) | 2 (4.8) | 0 (0.0) |
| HR negative |  |  |  |  |
| grade 1 | 0 (0.0) | 0 (0.0) | 0 (0.0) | 0 (0.0) |
| grade 2 | 9 (8.0) | 5 (17.9) | 1 (2.4) | 3 (7.1) |
| grade 3 | 37 (33.0) | 12 (42.8) | 9 (21.4) | 16 (38.1) |
| unknown | 1 (0.9) | 1 (3.6) | 0 (0.0) | 0 (0.0) |
| **average number of pre-operative chemotherapy cycles (range)** | 5.3 (2-8) | 5.1 (2-8) | 5.1 (2-6) | 5.5 (3-8) |
| **number of pre-operative chemotherapy cycles** |  |  |  |  |
| ≤4 | 34 (30.4) | 9 (32.1) | 15 (35.7) | 10 (23.8) |
| >4 | 78 (69.6) | 19 (67.9) | 27 (64.3) | 32 (76.2) |
| **chemotherapy regimens** |  |  |  |  |
| carboplatin + docetaxel or paclitaxel | 21 (18.8) | 21 (75.0) | - | - |
| cisplatin + docetaxel or paclitaxel | 7 (6.3) | 7 (25.0) | - | - |
| AC | 2 (1.8) | - | 2 (4.8) | - |
| FEC | 15 (13.4) | - | 15 (35.7) | - |
| CMF | 1 (0.9) | - | 1 (2.4) | - |
| AC + docetaxel or FEC + docetaxel | 24 (21.4) | - | 24 (57.1) | - |
| epirubicin + docetaxel or paclitaxel | 32 (28.5) | - | - | 32 (76.2) |
| doxorubicin + docetaxel or paclitaxel | 10 (8.9) | - | - | 10 (23.8) |
| **pre-operative trastuzumab** | 15 (13.4) | 8 (28.6) | 5 (11.9) | 2 (4.8) |
| **ypT** |  |  |  |  |
| <2cm | 42 (37.5) | 11 (39.3) | 17 (40.5) | 14 (33.3) |
| ≥2cm | 69 (61.6) | 16 (57.1) | 25 (59.5) | 28 (66.7) |
| unknown | 1 (0.9) | 1 (3.6) | 0 (0.0) | 0 (0.0) |
| **ypN** |  |  |  |  |
| negative | 31 (27.7) | 9 (32.1) | 11 (26.2) | 11 (26.2) |
| positive | 75 (67.0) | 18 (64.3) | 29 (69.0) | 28 (66.7) |
| unknown | 6 (5.3) | 1 (3.6) | 2 (4.8) | 3 (7.1) |
| **ΔStrTIL** |  |  |  |  |
| zero or positive | 87 (77.7) | 22 (78.6) | 35 (83.3) | 30 (71.4) |
| negative | 25 (22.3) | 6 (21.4) | 7 (16.7) | 12 (28.6) |

AC: doxorubicin plus cyclophosphamide, CMF: cyclophosphamide plus methotrexate plus 5-fluorouracil, ILC: invasive lobular carcinoma, FEC: 5-fluorouracil plus epirubicin plus cyclophosphamide, HR: hormone receptor, NST: no special type, StrTIL: stromal tumor-infiltrating lymphocytes

According to the type of pre-operative chemotherapy, the patients were grouped into platinum-based, cyclophosphamide-based and anthracycline-based groups. The treatment regimens are presented in Table 1. All treatment regimens were of conventional doses and schedules, and selected based on valid international guidelines.

### Pathology

Formalin fixed, paraffin embedded blocks of core biopsies and surgical specimens were retrieved from the four pathology departments.

4μm sections of representative tumor blocks were stained with hematoxylin and eosin (H&E). The percentage of stromal tumor-infiltrating lymphocytes (StrTIL) was evaluated according to the recommendation of International TILs Working Group 2014 [10]. Histopathologic evaluation of StrTILs was performed by GCs, AMT, AV, ET and JK. Controversial cases were reevaluated and discussed.

### Statistical analyses

SPSS 20.0 (SPSS Inc., Chicago, IL, USA) was used for statistical analysis. The normality of the data was controlled by the Shapiro-Wilk test. The association between changes in StrTIL and clinico-pathological variables (pre-operative chemotherapy received, grade, immunohistochemical subtype and age) was calculated by the Wilcoxon-signed rank test.

The distant metastasis-free survival (DMFS) was assessed and defined as the time interval between the first cycle of pre-operative chemotherapy and the date of distant relapse or death. Nine cases with bone metastases at baseline were censored from the DMFS analyses. Database was locked in December 2017. The prognostic value of StrTIL changes (ΔStrTIL: the difference between post-treatment stromal tumor-infiltrating lymphocytes in surgical specimen (post-StrTIL) and pre-treatment stromal tumor-infiltrating lymphocytes in core biopsy (pre-StrTIL) was tested as continuous variable. Hazard ratios and 95% confidence intervals (95% CI) were calculated with the Cox proportional hazard regression model. Multivariate Cox regression analysis included the following prognostic factors: age, grade, HR status, type of treatment, residual tumor size and post-treatment pathological lymph node status. The Kaplan–Meier method (the log-rank test) was used to analyze the role of ΔStrTIL in DMFS in HR negative and HR positive tumor groups separately.

All applied statistical tests were two-sided.

## Results

### Baseline disease characteristics

Samples from 112 individuals were available for analysis. Most patients (86.6%, n=97) had invasive carcinoma of no special type (NST), 42.0% (n=47) were HR negative and 58.0% (n=65) were HR positive. The clinico-pathological characteristics are reported in Table 1. At initiation of pre-operative chemotherapy the patients had a mean age of 55 years (range: 29- 80 years). 25.0% of the patients (n=28) received platinum-based therapy, 37.5% (n=42) received cyclophosphamide-based therapy and 37.5% (n=42) received anthracycline-based therapy.

Of the 28 patients undergoing platinum-based therapy, 64.3% (n=18) were HR negative (mainly triple negative, 46.4% (n=13)) and 35.7% (n=10) were HR positive. According to the chemotherapy regimen used, 21 patients were treated with carboplatin + docetaxel or paclitaxel and 7 patients received cisplatin + docetaxel or paclitaxel. Our previously published data suggested that the majority of mutations are produced by platinum in this cohort and the expected number of neo-epitopes is about 20-fold higher than in a cohort treated only with anthracycline and/or taxane [5].

Of the 42 patients undergoing cyclophosphamide-based therapy, 23.8% (n=10) were HR negative and 76.2% (n=32) were HR positive. 57.1% (n=24) received anthracycline (epirubicin (E) or doxorubicin) + cyclophosphamide (C) in combination with or without 5-fluorouracil (F) followed by docetaxel. The other commonly used treatment regimen in this group was FEC without following taxane therapy in 35.7% (n=15) of the cases (Table 1). Our previously published data suggested that the majority of mutations are produced by cyclophosphamide in this cohort and the expected number of neo-epitopes is about 5-fold higher than in a cohort treated only with anthracycline and/or taxane [5].

Of the 42 patients undergoing anthracycline-based therapy, 45.2% (n=19) had HR negative and 54.8% (n=23) had HR positive carcinomas. All cases were treated with anthracycline + taxane combination (of those 32/42 received epirubicin + docetaxel or paclitaxel).

The majority of patients received more than four cycles of chemotherapy and the average cycle number was the same in each group (Table 1).

Of the 22 HER2 positive cases, 68.2% (n=15) received pre-operative trastuzumab therapy. Trastuzumab was administered in combination with platinum-based (n=8), cyclophosphamide-based (n=5) or anthracycline-based (n=2) therapy (Table 1).

### StrTIL changes before and after chemotherapy

In the pre-treatment core biopsy samples, the median pre-StrTIL was 3.00% (interquartile range (IQR): 1.00-7.50) and more than 50% StrTIL (lymphocyte predominant) was detected in only one case. The post-StrTIL reached 50% or above in 10 cases (the pre-operative therapy was platinum-based (n=4), FEC (n=1) or docetaxel + epirubicin (n=5)), (Fig. 1a-i). The median post-StrTIL rose significantly to 6.25% (IQR: 3.00-25.00; p<0.001) after treatment. Pre-StrTIL less than 1% was observed in 14 cases, while StrTIL less than 1% in the residual tumor occurred in only two cases.

**Figure 1.**
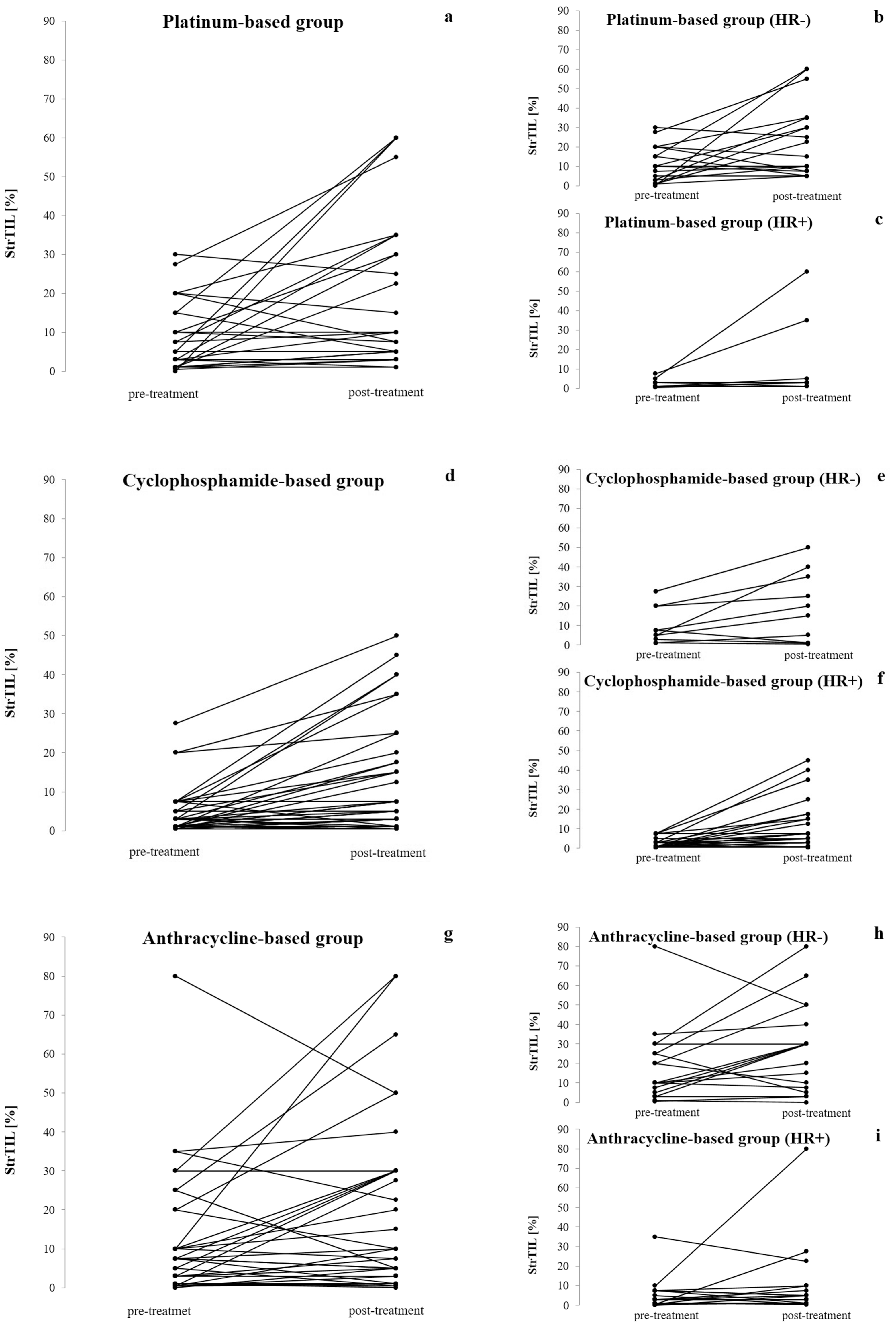
Stromal tumor-infiltrating lymphocytes (StrTIL) before and after pre-operative chemotherapy. Significant ΔStrTIL increase was observed in the three treatment groups (platinum-based: p=0.007; cyclophosphamide-based: p<0.001; anthracycline-based: p=0.047; Fig. 1a, d, g). By analyzing separately the HR positive and HR negative cases we experienced only the administration of cyclophosphamide resulted in significant ΔStrTIL increment in HR positive cases (p<0.001, Fig. 1c, f, i), whereas in HR negative cases, a prominently strong relationship between the treatment applied and StrTIL changes could not be proven (platinum-based: p=0.026; cyclophosphamide-based: p=0.049; anthracycline-based: p=0.063).

The increase in post-StrTIL was significant both in HR positive (ΔStrTIL positive: n=32 (49.2%); zero: n=21 (32.3%); negative: 12 (18.5%)) and HR negative (ΔStrTIL positive: n=29 (61.7%); zero: n=5 (10.6%); negative: n=13 (27.7%)) cases (p<0.001 in both groups; Table 2). In the subgroup of HR positive/HER2 negative cases, the changes in StrTIL was significant in grade 3 cases (ΔStrTIL positive: n=14 (66.7%); zero: n=3 (14.3%); negative: n=4 (19.0%); p=0.007) but not in grade 1-2 cases (ΔStrTIL positive: n=11 (36.6%); zero: n=14 (46.7%); negative: n=5 (16.7%); p=0.075; Table 2).

**Table 2.**
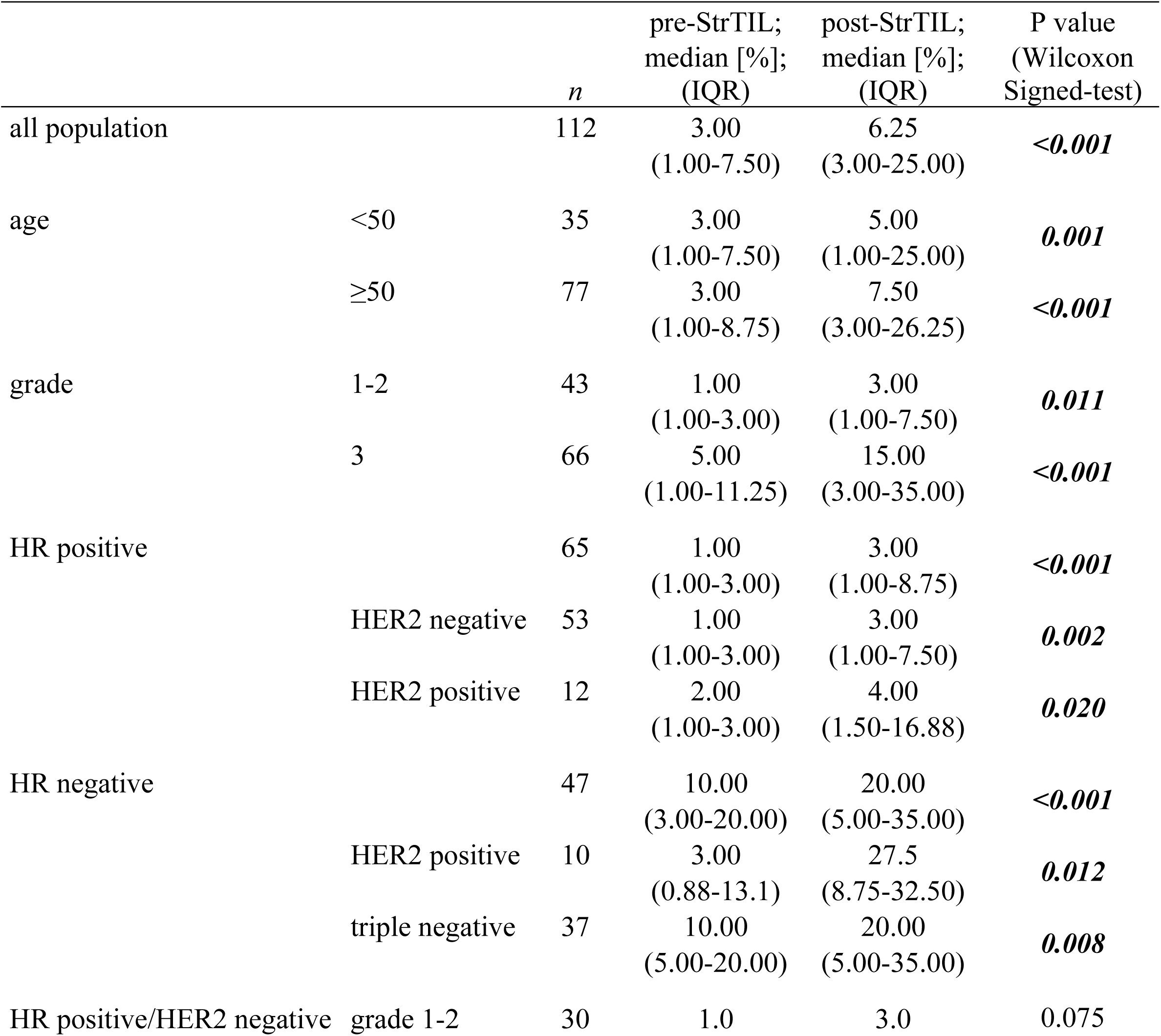
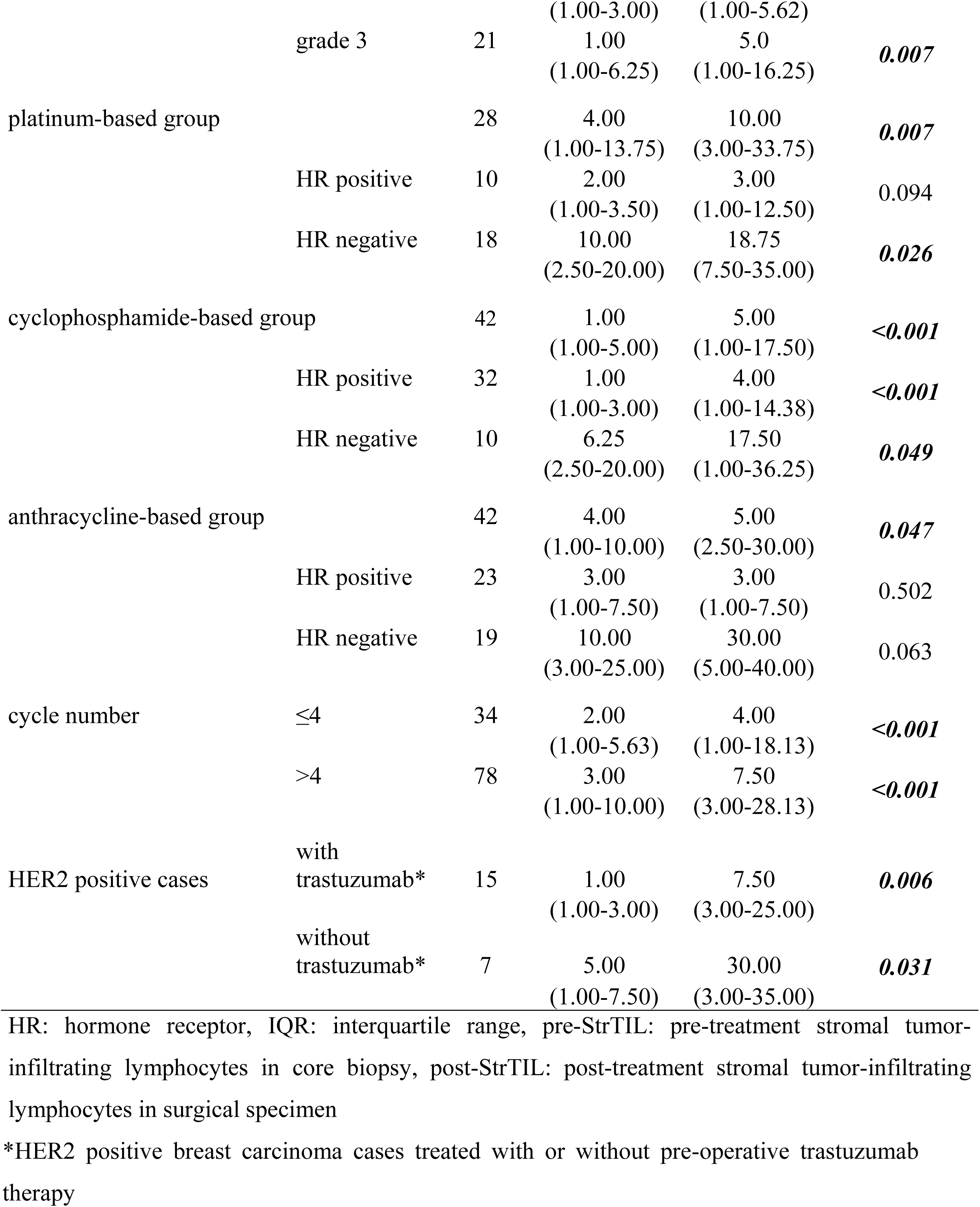
Changes in stromal tumor-infiltrating lymphocytes (StrTIL): median StrTIL levels before and after pre-operative chemotherapy

We did not detect any association between changes in StrTIL and other features (shown in Table 2).

When analyzing the pre-StrTIL and post-StrTIL among the three treatment groups, we experienced significant StrTIL increase independently from the treatment applied (Table 2; Fig. 1a, 1d, 1g; Supplementary Table 1). Interestingly, in the subgroup analysis, only the administration of cyclophosphamide resulted in a significant increase in StrTIL in HR positive cases (ΔStrTIL positive: n=18 (56.3%); zero: n=10 (31.2%); negative: n=4 (12.5%); p<0.001; Table 2; Fig. 1c, 1f, 1i; Supplementary Table 1).

### Survival analyses

Data on DMFS was available for 103 cases. The median DMFS was 28.2 months (range: 2.6-118.3 months). Distant metastases occurred in 31/103 (30.1%) cases. In 21/31 (67.7%) cases, the primary breast carcinoma was HR negative, and in 19/31 (61.3%) cases the post-StrTIL was lower than 10.0% or showed a decrease in comparison with the pre-StrTIL. As reported in Table 3, in univariate analyses, the HR status and the post-treatment pathological lymph node status were the only significant factors influencing DMFS. In the multivariate model changes of StrTIL showed a strong prognostic value (Table 3). The Cox analysis in HR negative cases confirmed both post-StrTIL and ΔStrTIL as playing independent prognostic role in DMFS. Each 1% increase in post-StrTIL reduced the hazard of distant metastases development by 2.6% (hazard ratio: 0.974; CI: 0.948-1.000; p=0.05) and for each 1% ΔStrTIL increment, the risk of distant metastases was reduced by 4.3% (hazard ratio: 0.957; CI: 0932-0.983; p=0.001), but according to our results, the pre-StrTIL did not influence the DMFS. The prognostic role of StrTIL in HR positive cases could not be proven (Supplementary Table 2). The Kaplan-Meier analysis was carried out in HR negative and HR positive cases separately. Among HR negative cases, increased or unchanged post-StrTIL was associated with better survival (Fig. 2c).

**Table 3.**
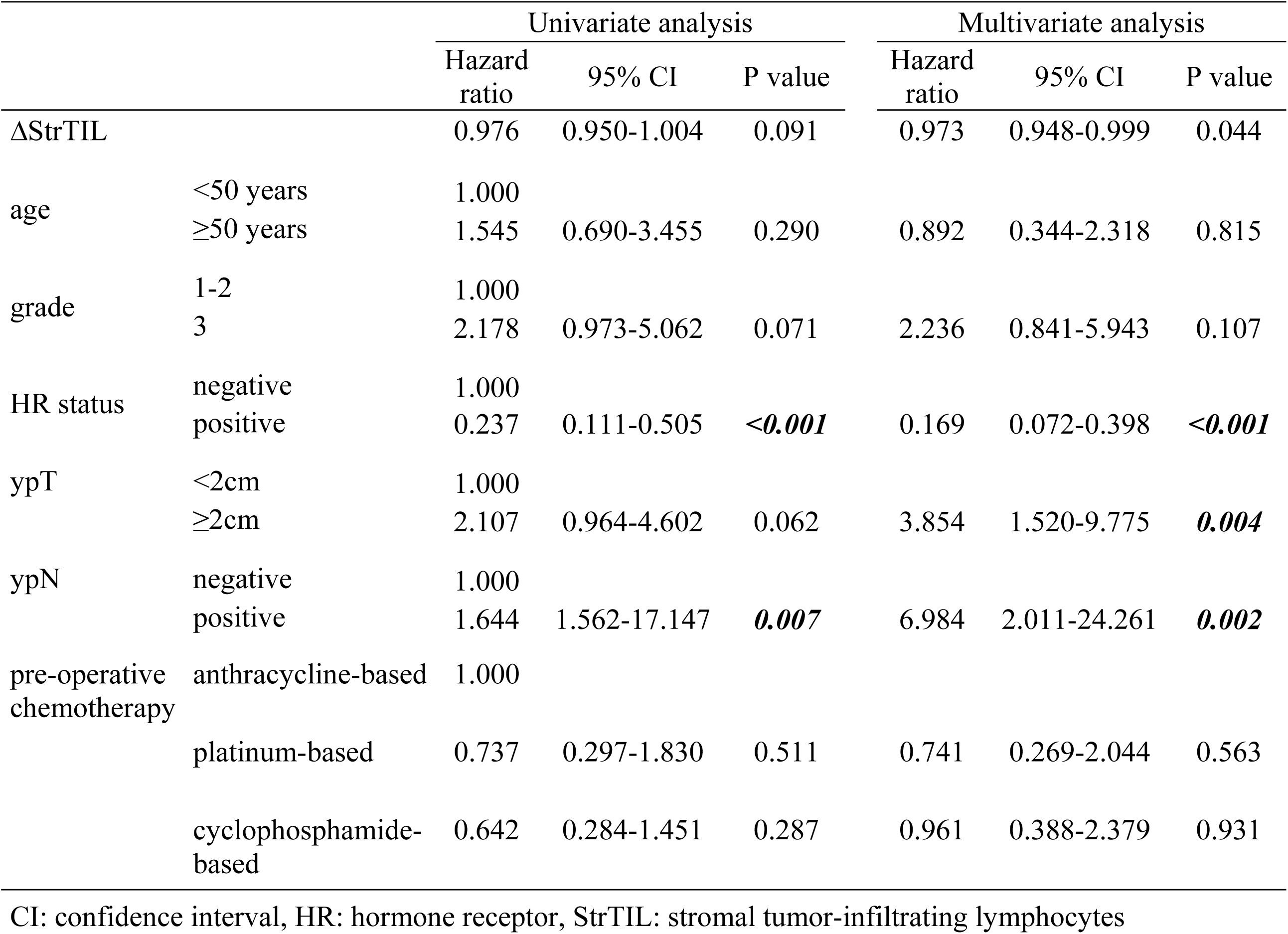
Factors associated with distant metastasis-free survival

**Figure 2.**
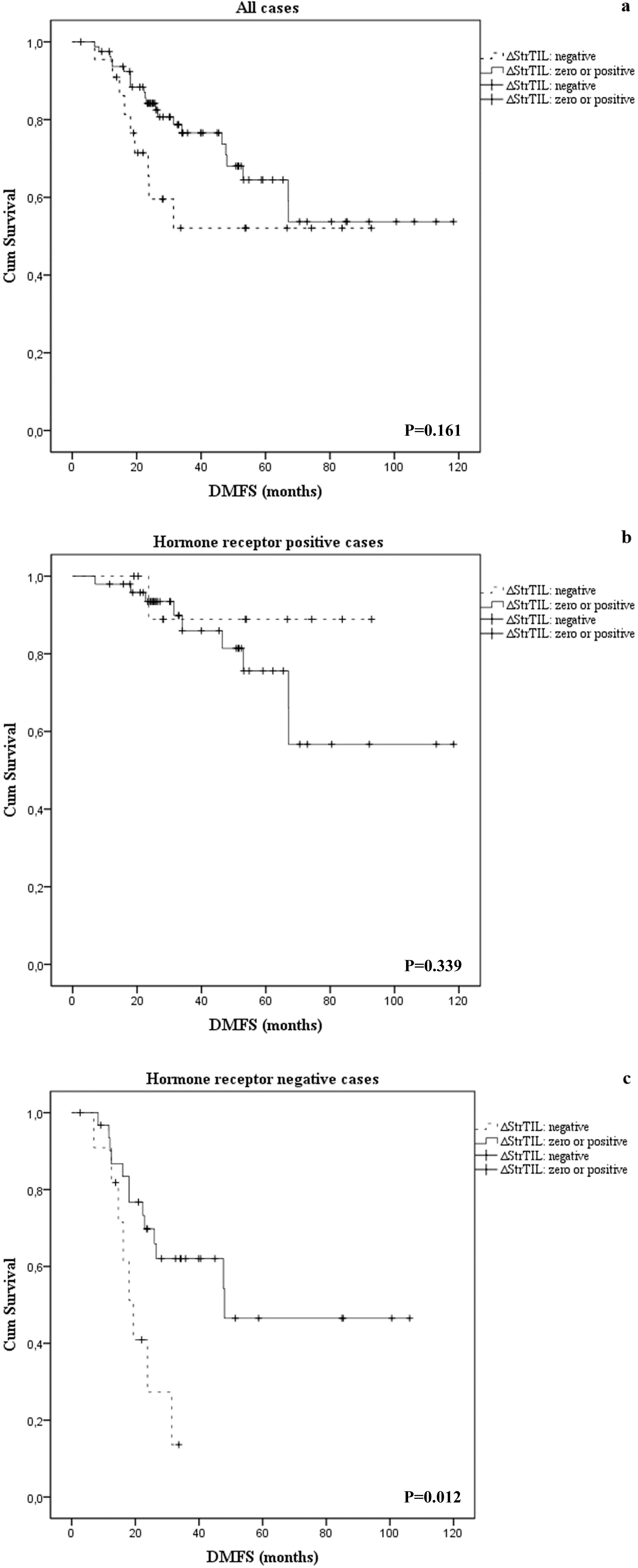
Kaplan-Meier curves of survival analyses. By analyzing the whole study cohort no significant correlation was detected between ΔStrTIL and distant metastasis-free survival (DMFS) (p=0.161, Fig. 2a). The same result was observed in hormone receptor positive cases too (p=0.339, Fig. 2b). In hormone receptor negative cases the estimated median DMFS was significantly higher, if the ΔStrTIL was zero or positive (48.0 months; standard error: 8.7) compared to the cases where ΔStrTIL was negative (19.4 months; standard error: 2.4) (Fig. 2c).

## Discussion

It has been established in several studies that tumor infiltrating lymphocytes in baseline, pre-treatment biopsies of breast cancer are powerful prognostic markers of the clinical outcome in triple negative and HER2 positive breast cancers [11,12]. The correlation between post-treatment levels of TIL and clinical outcome is less clear, perhaps due to the fact that cases with pathological complete response are often eliminated from further analysis. High post-treatment TIL levels were associated with better outcome in some studies [13,14] but others did not find similar correlations [15,16]. Most recently, the prognostic role of change in StrTIL levels between the pre- and post-treatment samples was investigated and an increase in StrTIL levels was associated with better survival [15]. Here we are confirming this observation, suggesting that the increase of TIL ratio in residual disease as a surrogate measure of anti-tumor immune activation may in fact reflect significant therapeutic benefit. Cytotoxic chemotherapy has been shown to increase T cell response in breast cancer [13,15]. The various agents may improve immune response in vivo by a wide array of biological mechanisms. Previous studies have shown that anthracyclines, such as doxorubicin may induce T cell activation in breast cancer by a toll-like receptor driven mechanism [17], while cyclophosphamide may enhance anti-tumorigenic immune response by suppressing T regulatory cell function [18]. Importantly, it was shown in preclinical studies that artificial inactivation of MutL homologue 1 (MLH1) increased the mutational burden and led to dynamic mutational profiles, which resulted in the persistent renewal of neo-antigens in vitro and in vivo, improving immune surveillance [19]. Therefore, we hypothesized that induction of somatic tumor mutations via mutagenic chemotherapy would increase the immunogenicity of tumors and improve immune responses in patients with breast cancer. Furthermore, if a cytotoxic chemotherapy agent induces significantly more mutations/neo-epitopes it is also expected to induce a more intense immune reaction, as reflected in the number of TILs. Such a correlation would justify the combination of highly mutagenic agents with immune checkpoint inhibitor therapy [20]. This was the rationale behind comparing the TIL induction by three classes of cytotoxic agents, each with a distinct level (high, medium and low) mutagenic capacity. Platinum showed a 20-fold higher and cyclophosphamide showed a 5-fold higher mutagenic and probably neo-epitope inducing capacity than anthracycline or taxane therapy [5]. Previous studies did not attempt to perform such a comparison and in particular no data were reported on the induction of TILs by platinum-based neoadjuvant chemotherapy in breast cancer.

Based on our previous findings [5], we expected significantly more TIL induction by the highly mutagenic platinum agent relative to treatments without platinum or cyclophosphamide. However, while platinum and cyclophosphamide-based treatment seemed to have induced a somewhat more significant increase in TIL, neoadjuvant chemotherapy containing none of these agents also increased post-treatment TIL. This does not support the notion that chemotherapeutic agents showing 5-20 fold higher mutagenic potential in experimental systems would be better candidates to be combined with immune checkpoint inhibitors.

There are several possible explanations for this lack of difference. First, the mutagenicity of the therapeutic agents was tested in cell lines, and currently there are no reliable measures of mutagenic capacity of these agents in human tumor samples. Therefore, we do not know whether the difference seen in cell lines also holds in human tumors in vivo. It is also possible that the higher mutagenicity of a given agent, e.g. platinum, is compensated by another mechanism, such as the downregulation of the MHC (major histocompatibility complex) complex [21]. While this is possible, it should be noted that platinum treatment was reported to induce HLA (human leukocyte antigen) expression in breast cancer [22]. Finally, it was suggested that in breast cancer, the main increase in therapeutic TIL response is driven by gamma delta lymphocytes and not alpha beta lymphocytes, and the former response is typically not induced by neo-epitopes [23].

## Conclusion

In summary, we did not find a significantly higher level of StrTIL induction by more mutagenic chemotherapeutic agents. Therefore, combining those with immune checkpoint inhibitors may not lead to enhanced therapeutic benefit.

## List of abbreviations

AC: doxorubicin plus cyclophosphamide
CI: confidence interval
CMF: cyclophosphamide plus methotrexate plus 5-fluorouracyl
DMFS: distant metastasis-free survival
FEC: 5-fluorouracyl plus epirubicin plus cyclophosphamide
HER2: human epidermal growth factor receptor 2
HLA: human leukocyte antigen
HR: hormone receptor
MHC: major histocompatibility complex
MLH1: MutL homologue 1
ILC: invasive lobular carcinoma invasive carcinoma
NST: invasive carcinoma of no special type
IQR: interquartile range
post-StrTIL: post-treatment stromal tumor-infiltrating lymphocytes in surgical specimen
pre-StrTIL: pre-treatment stromal tumor-infiltrating lymphocytes in core biopsy
StrTIL: stromal tumor-infiltrating lymphocytes

## Declarations

### Ethics approval and consent to participate

The study was approved by the Hungarian Medical Research Council (ETT-TUKEB 14383/2017) and it was conducted in accordance with the Declaration of Helsinki.

### Consent for publication

Not applicable.

### Availability of data and material

The datasets used and/or analyzed during the current study are available from the corresponding author on reasonable request.

### Funding

New National Excellence Program (ÚNKP-17-4-II-SE-65) (L.V)

New National Excellence Program (ÚNKP-17-4-III-SE-71) (AMT)

NVKP_16-1-2016-0004

STIA 19/2017, 6800313113, 68003F0043 (AMT)

The Research and Technology Innovation Fund (KTIA_NAP_13-2014-0021 to Z.S.)

NKP-2017-00002 (Z.S.)

Breast Cancer Research Foundation (Z.S.)

### Authors’ contributions

AMT performed the histopathologic evaluation of stromal tumor-infiltrating lymphocytes, participated in acquisition, interpretation of data and was involved drafting the manuscript.

OR made substantial contributions to design, acquisition of data, analyses and interpretation of data and was involved drafting the manuscript.

GCS performed the histopathologic evaluation of stromal tumor-infiltrating lymphocytes, took part in acquisition of data and revised the manuscript critically for important intellectual content.

ET performed the histopathologic evaluation of stromal tumor-infiltrating lymphocytes, revised the manuscript critically for important intellectual content.

GR participated in data collection, read and approved the final manuscript.

TT participated in data collection, revised the manuscript critically for important intellectual content.

LV provided professional advices for statistical analyses and revised the manuscript.

LR contributed to the research by constructive criticism and revised the final manuscript.

RK participated in data collection, read and approved the final manuscript.

ZSK revised the manuscript critically for important intellectual content and approved the final version.

JK performed the histopathologic evaluation of stromal tumor-infiltrating lymphocytes, revised the manuscript and approved the final version.

MD contributed to the research by constructive criticism, read and approved the final manuscript.

AV performed the histopathologic evaluation of stromal tumor-infiltrating lymphocytes, read and accepted the final manuscript.

ZSZ made substantial contributions to conception, was involved in drafting the manuscript and revised it critically for important intellectual content, read and approved the final manuscript.

## Acknowledgements

Not applicable.

## Suppl. Table 1..xlsx

**Supplementary Table 1. Changes of stromal tumor-infiltrating lymphocytes in the three treatment groups**

HR: hormone receptor, StrTIL: stromal tumor-infiltrating lymphocytes HR: hormone receptor, StrTIL: stromal tumor-infiltrating lymphocytes

## Suppl. Table 2..xlsx

**Supplementary Table 2. Prognostic value of pre-treatment StrTIL, post-treatment StrTIL and changes in StrTIL**

CI: confidence interval, pre-StrTIL: pre-treatment stromal tumor-infiltrating lymphocytes, post-StrTIL: post-treatment stromal tumor-infiltrating lymphocytes

